# scXOrGAN: A Generative Framework for Cross-Organ Single-Cell Transcriptomic Translation

**DOI:** 10.64898/2026.11.25.769347

**Authors:** Farriha Afnan, Ananya Shahrin Promi, Md. Abul Hassan Samee, Atif Rahman

## Abstract

Gene expression profiles of organs provide important insights into their health and disease states. However, while some organs can be sampled only through invasive procedures, others, such as blood, are readily accessible. Existing methods for predicting gene expression in one organ from another typically work with bulk or averaged profiles, obscuring cell specific differences. Translation at single-cell resolution is more challenging, because individual cells in the source and target organs lack direct correspondence. Here, we present scXOrGAN, a generative model that translates single-cell transcriptomes between two organs of the same individual using data that are unpaired at the cell level. scXOrGAN combines two variational autoencoders that share a latent space with cycle consistency, adversarial training and distribution matching. Across three organ pairs (ileum to rectum, blood to bronchoalveolar lavage fluid (BALF) in COVID-19, and blood to heart in heart failure), the generated gene expressions closely matched the real target cell profiles of held-out donors and preserved cell-to-cell variation. Notably, generated BALF macrophages exhibit expression of COVID-19-associated genes that are nearly absent in source blood cells, while generated cardiac fibroblasts express genes upregulated in heart failure.

## INTRODUCTION

Although all organs of an individual share the same genome, the set of genes expressed varies from one organ to another. Cross-organ translation refers to predicting the transcriptome of one organ from that of another organ in the same individual. As organs are functionally interconnected, are perfused by a common circulation and communicate through cell signaling, physiological and pathological states in one organ are reflected in others^1,2^. Blood in particular has previously been observed to reflect physiological and pathological changes elsewhere in the body^3^. Clinical practice already exploits this principle, for example through blood-based markers of liver and kidney function. Cross-organ translation extends it from individual markers to the whole transcriptome.

Cross-organ translation has the greatest potential in the diagnosis of diseases. Every organ has its own molecular signature^4,5^ that is altered by diseases. However, organs differ widely in how readily they can be sampled. Some tissues can be accessed only through invasive procedures that carry substantial risk, while others such as blood are collected routinely and with minimal harm. This raises the question of whether the transcriptome of an inaccessible tissue can be inferred from that of an accessible one. If so, the molecular state of a hard-to-reach organ could be monitored more frequently and at lower risk to the patient, with potential applications ranging from disease progression and treatment response to the surveillance of transplanted organs.

Cross-organ translation has already been shown at the bulk level. TEEBoT showed that the whole blood transcriptome can predict tissue-specific expression for approximately 60% of genes on average^6^, and MTM later improved on this with a deep multi-task learning framework^7^. TEEBoT also showed that the predicted tissue expression could predict disease state for six diseases, such as hypertension and type 2 diabetes, almost as well as the real tissue expression^6^. However, these methods operate on bulk profiles, where cell-type specific signal is lost. A bulk profile is an average over all the cells in a sample, so a change that happens in only one cell type can disappear in the average. TIMEP took a step towards single-cell resolution, predicting the immune cells of head and neck tumors from matched blood single-cell data^8^. But it first averages each cell type into a single pseudo-bulk profile and its output is still one average value per gene, not a population of cells.

We ask the same question at single-cell resolution, and this brings a new challenge. While we can assay cells from both organs of the same person, there is no correspondence between any two single cells. The unavailability of paired source organ to target organ data at the cell level motivates us to adopt an approach similar to CycleGAN^9^, a deep learning framework for unpaired image-to-image translation, where cycle-consistency substitutes for missing pairwise supervision between two unpaired populations.

Similar adversarial, cycle-consistent ideas have been used at single-cell resolution, but for cross-omic translation within a single tissue. scACT extracts cross-modality biological insights from unpaired single-cell data using adversarial training^10^; scMMGAN aligns modalities such as scRNA-seq and scATAC-seq measured on different cells from the same population^11^; and UnCOT-AD translates between unpaired omics layers for Alzheimer’s disease prediction^12^. Single-cell perturbation models such as scGen^13^ and scPRAM^14^ also learn from unpaired cells, but focus on predicting cellular responses to perturbations rather than translating cells across organs. None of these methods cross the organ or tissue boundary itself.

To address the challenge of non-invasive, disease relevant monitoring of hard-to-sample organs at single-cell resolution, we propose scXOrGAN, a cycle-consistent dual-VAE and adversarial framework that translates between a source organ and a target organ at single-cell resolution. Because the output remains single-cell resolved, our approach yields cell-type specific distributions rather than a single averaged estimate per gene. We test scXOrGAN on datasets from three pairs of organs: ileum to rectum^15^, blood to BALF in COVID-19^16^, and blood to heart in heart failure^17^. We compare it with scGen and scPRAM, two widely used methods for the closest related task.

## RESULTS

### Overview of scXOrGAN

Figure 1A illustrates the motivation of scXOrGAN. Some organs, such as blood, are easy and safe to sample. Others, such as the heart, can only be sampled through invasive and risky procedures. For a patient, we can measure the gene expression of single cells from the easy-to-access organ, which we call the source organ. We want to generate the gene expression of cells from the hard-to-reach organ, which we call the target organ.

**Figure 1.**
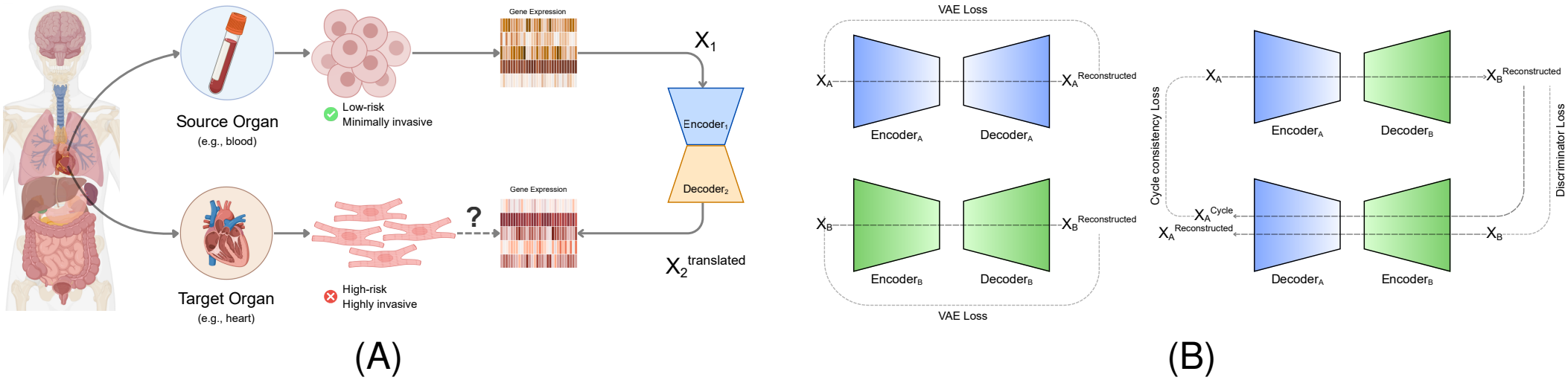
Overview of scXOrGAN. (A) The cross-organ translation problem. Single cells can be collected from a source organ that is easy and safe to sample, such as blood, whereas cells of a target organ, such as the heart, can be obtained only through invasive, high-risk procedures. Given the gene expression profiles of a patient’s source-organ cells (*X*_1_), scXOrGAN encodes each cell with the source-organ encoder and decodes it with the target-organ decoder, generating target-organ cells of the same patient 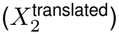. (B) Architectural overview of scXOrGAN. Left: two variational autoencoders (VAEs), one for the source organ (blue) and one for the target organ (green), each reconstruct their own input and are trained with the VAE loss (reconstruction error and KL divergence). Both encoders map cells into one shared latent space. Right: translation and cycle consistency. A source-organ cell *X*_*A*_ is encoded by Encoder_*A*_ and decoded by Decoder_*B*_ into a translated target-organ cell, which is then encoded by Encoder_*B*_ and decoded by Decoder_*A*_ back into the source organ 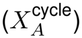. The cycle-consistency loss penalizes the difference between *X*_*A*_ and 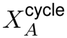; the same is done in the direction from B to A. A discriminator compares translated cells with real cells of the target organ (discriminator loss). The latent discriminator and the distribution-matching (MMD) term are not drawn; all losses are defined in Methods.

The main challenge is that there is no one-to-one correspondence between cells of the two organs. What we do have is cells from both organs of the same patients. So the training data are matched at the patient level, but unpaired at the cell level. This rules out ordinary supervised learning, where each input comes with its correct output.

To learn from such data, scXOrGAN uses an architecture similar to the one deployed by Cy-cleGAN^9^ in the context of images. In our adaptation, an encoder compresses the expression of a cell into a short summary. A decoder then turns such a summary back into gene expression. If cells from both organs are summarized in the same space, then a summary made from a source cell can be decoded as a target cell^18^. scXOrGAN is built around a dual-branch variational autoencoder (Dual-VAE)^19^, with one branch for the source organ and one for the target organ, as illustrated in Figure 1B. Each branch encodes gene expression profiles into a shared latent space via a variational encoder, and decodes back into expression space via a decoder.

Cross-organ translation is achieved by cross-wiring the branches: a source-organ profile is encoded by the source encoder and decoded by the target decoder to produce a translated target profile. Since there is no true matching cell, we cannot check a translated cell directly. Instead, cycle-consistency loss^9^ ensures a profile translated to the other domain and back recovers the original; so the model cannot just learn any mapping that satisfies the discriminators, it has to learn one that keeps each translated cell tied to the cell it originated from.

The model is guided toward realistic outputs through adversarial discriminators^20^: one aligning the shared latent space across domains, and one per domain distinguishing real from generated expression profiles. Discriminators judge single cells, so we also compare whole groups of cells. A distribution-matching term, the maximum mean discrepancy (MMD)^21^, checks whether the generated cells and the real target cells of a patient have the same overall distribution. Source and target-organ cells are matched at the patient level during training, while remaining unpaired at the cell level; no individual source cell is assumed to correspond to any individual target cell. We use this patient-level matching directly: in each training step, all cells come from a single patient, so generated cells are always compared with real cells of the same person. Once trained, scXOrGAN needs only the source-organ cells of a new patient. Each source cell is translated into one target cell, so the output is a population of cells rather than a single average profile. Details of the model and its training are given in Methods.

Testing scXOrGAN requires single-cell RNA-seq data from two organs of the same donors, and few public datasets meet this requirement. Many of those that do profile only immune cells in both organs. The immune cell types are highly correlated between organs and are nearly identical in expression, so translating them from one organ to the other would not provide a meaningful test of the model. We therefore selected datasets that also profile cell types of the target organ whose expression differs substantially from that of the source organ’s cells. We identified three datasets to meet these criteria.

The first is IBDverse^15^, which contains samples of the terminal ileum and the rectum from healthy donors. Its large number of donors sampled in both organs provides a larger set of held-out donors for testing. From this dataset, we translated two cell types in the ileum, memory B cells and CD8 memory T cells, into colonocytes in the rectum. The second is a COVID-19 atlas^16^ with blood and bronchoalveolar lavage fluid (BALF) from the same patients, which links the two organs to a disease. Here we translated four cell types in blood into macrophages in BALF, and CD8 T cells in blood into epithelial cells in BALF. Macrophages in BALF are immune cells, but they are a different cell type from the blood cells used as the source and differ considerably from them in expression. The third is a heart failure atlas^17^ with blood and heart tissue from the same patients, which also links the two organs to a disease. Here we translated B cells and T cells in blood into endothelial cells and fibroblasts in the heart.

For each dataset, we trained and tested scXOrGAN in the same way. We trained one model for each pair of cell types, for example CD8 T cells in blood to macrophages in BALF. The donors of each dataset were split into training and test donors, with all data from test donors excluded from model training. At test time, it received only the source-organ single-cell RNA-seq(scRNA-seq) data of a test donor and generated the target-organ cell profiles, which we then compared with the corresponding real target-organ cell data from the same donor, withheld during training. All genes used by the model were chosen from the training donors only.

Because the two organs have no matching cells, we cannot check a generated cell against its corresponding cell. So we evaluate the generated cells as a group. First, we compare the average expression of each gene in the generated and the real cells, which tells us whether the model produces the right type of cell. Second, we compare distributions and whether the generated cells vary from cell to cell as real cells do, rather than all being copies of one average profile. For this we use the energy distance (E-distance)^22^, which measures how far apart two groups of cell profiles are and is widely used to compare single-cell populations^23^. Third, we check what is specific to each patient, by asking whether the model predicts how a test patient differs from the average of the training patients. Since no method has been developed for this exact task previously, we compare scXOrGAN with two methods adapted from the closest task, single-cell perturbation prediction: scGen^13^ and scPRAM^14^. We also use a simple baseline that assigns every generated cell the average target profile of the training patients.

### scXOrGAN translates expressions from ileal immune cells into rectal colon cytes

We first tested scXOrGAN on IBDverse^15^, which has biopsies of the terminal ileum and the rectum from the same healthy donors. We translated two cell types in the ileum, memory B cells and CD8 memory T cells, into colonocytes in the rectum. We used immune cells in the ileum as the source, as in the other two datasets where the source is blood, and colonocytes, the main epithelial cell type of the rectum, as the target. Among immune cell types, we took the B-cell and T-cell types with the most donors who had enough cells in both organs (at least 100 cells; 91 donors for memory B cells and 79 for CD8 memory T cells; Supplementary Data 1). For memory B cells in the ileum to colonocytes in the rectum, 76 donors were used for training and 15 were held out for testing. For CD8 memory T cells in the ileum to colonocytes in the rectum, 66 donors were used for training and 13 were held out.

Figure 2A shows, for one test donor, the average expression of each highly variable gene in real cells in the ileum (memory B cells or CD8 memory T cells) against the average expression in real colonocytes in the rectum of the same donor. Each point is a gene, and points on the dashed line have the same average in both. Before translation, the correlation was 0.49 for CD8 memory T cells in the ileum and 0.34 for memory B cells in the ileum, and many genes expressed in rectal colonocytes were close to zero in both ileal cell types. This gap shows that source cells cannot be used in place of target cells, because their average expressions are substantially different. Inferring the state of an organ from another therefore requires first translating the source cells into cells that resemble those of the target organ. Figure 2B shows the same comparison for the colonocyte profiles that scXOrGAN generated from the ileal cell scRNA-seq data. After translation, the points shifted toward the diagonal, with correlations of 0.98 and 0.97. This shows that the generated cells have the right cell identity, since giving every cell the average profile of rectal colonocytes from the training donors also reaches a similar correlation (0.98 and 0.97). Figures 2C and 2D show the distribution of per-gene average expression in real and generated colonocytes in the rectum for two test donors per pair. The generated distribution closely matched the real one for every donor.

**Figure 2:**
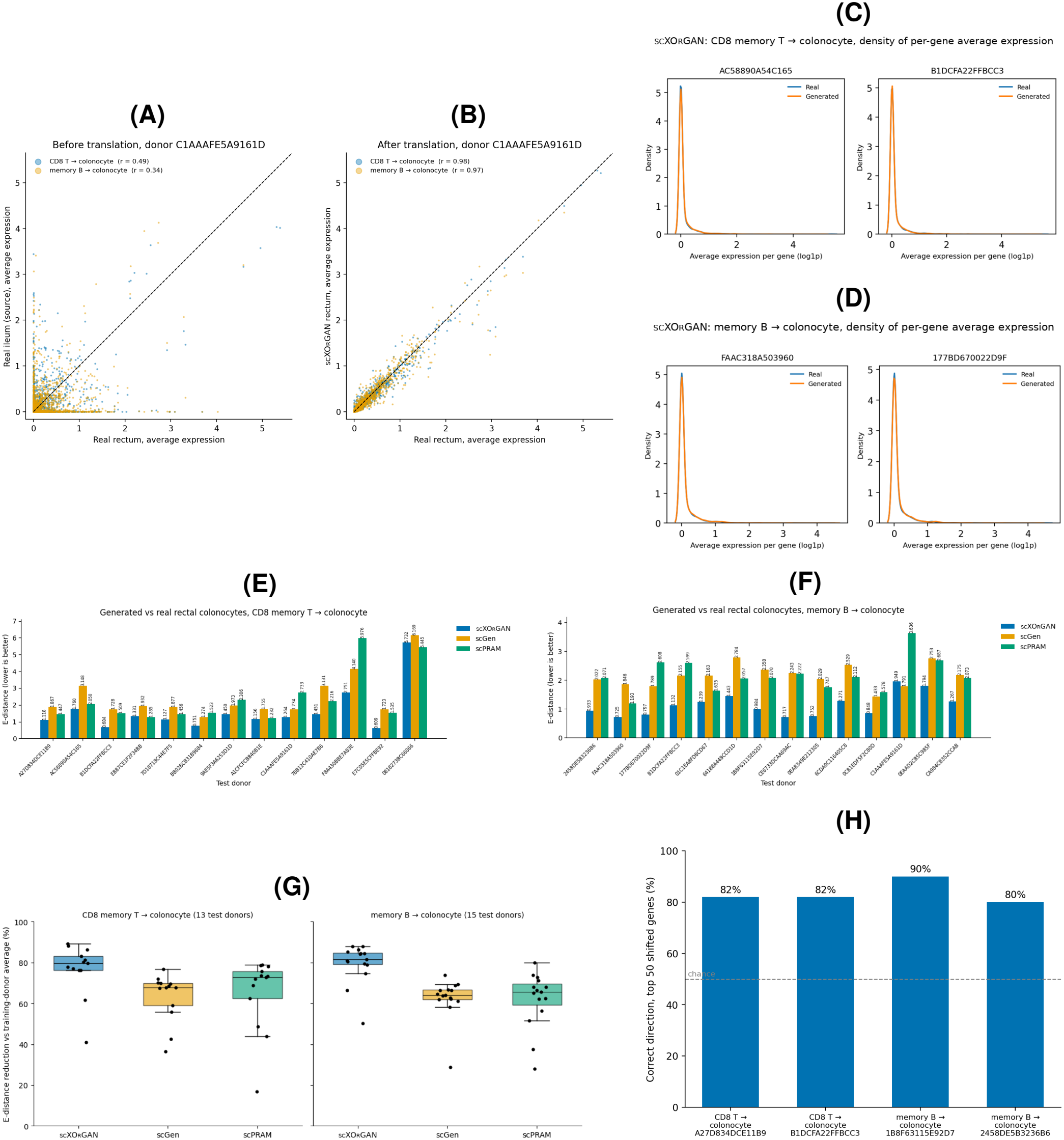
Translation of memory B cells and CD8 memory T cells in the ileum into colonocytes in the rectum by scXOrGAN. (A) Average expression of each highly variable gene in real source cells in the ileum (memory B cells or CD8 memory T cells) versus real colonocytes in the rectum, for test donor C1AAAFE5A9161D, colored by cell-type pair. (B) The same for scXOrGAN-generated colonocyte profiles versus real colonocytes in the rectum. *r*, Pearson correlation; dashed line, *y* = *x*. (C and D) Distribution of per-gene average expression in real and generated colonocytes in the rectum for two test donors, translated from CD8 memory T cells in the ileum (C) and memory B cells in the ileum (D). (E and F) E-distance between generated and real colonocytes in the rectum for each test donor and each method, translated from CD8 memory T cells in the ileum (E) and memory B cells in the ileum (F); lower is better. In (F), donor 0244A66A937BB2 is not shown because its values (7.2 to 10.5) are much larger than those of the other donors; it is included in (G). (G) E-distance reduction relative to the training-donor average (the average profile of rectal colonocytes in the training donors, assigned to every generated cell), for CD8 memory T cells in the ileum (13 test donors) and memory B cells in the ileum (15 test donors), both translated to colonocytes in the rectum. 0% is no better than the training-donor average. Boxes show median and interquartile range; points are individual donors. (H) Percentage of the 50 most-shifted genes in rectal colonocytes (largest real shift from the training-donor average) for which scXOrGAN predicted the correct direction, for the two test donors per pair with the highest score; dashed line, chance (50%).

We next compared the distribution of generated cells with that of real cells, rather than only their average expression. We trained scGen and scPRAM on the same donors, genes and number of epochs as scXOrGAN, and measured the E-distance between each method’s generated colonocyte profiles and that of real colonocytes in the rectum of each test donor. The E-distance is zero when two groups of cells have the same distribution, and grows as they differ. Figure 2F shows this E-distance for every test donor of the translation from memory B cells in the ileum to colonocytes in the rectum (for CD8 memory T cells in the ileum, see Figure 2E). scXOrGAN had a lower E-distance than scGen in 14 of 15 donors for ileal memory B cells and in all 13 donors for ileal CD8 memory T cells, and lower than scPRAM in all 15 and 11 of 13 donors. The median E-distance of scXOrGAN was 1.132 for ileal memory B cells and 1.264 for ileal CD8 memory T cells, against 2.163 and 1.877 for scGen and 2.073 and 1.535 for scPRAM. Because the scale of the E-distance differs between donors, we also measured each method’s reduction in E-distance relative to the training-donor average (Figure 2G). scXOrGAN reduced the E-distance by a median of 81.6% (range 50.1 to 87.9%) for ileal memory B cells and 79.6% (40.9 to 89.2%) for ileal CD8 memory T cells, compared with 64.0% and 67.8% for scGen and 65.6% and 72.9% for scPRAM. In 12 of 15 donors for ileal memory B cells and 10 of 13 donors for ileal CD8 memory T cells, the generated cell profiles were closer to the gene expressions of the real colonocytes of the donors in the rectum than that of real rectal colonocytes from the training donors were.

Finally, we checked whether scXOrGAN predicts the direction in which each donor differs from the training donors. For each test donor, we took the 50 genes with the largest real shift from the training-donor average in rectal colonocytes. Figure 2H shows, for the two test donors of each pair with the highest score, the percentage of these genes for which scXOrGAN predicted the correct direction. Across all test donors, scXOrGAN predicted the correct direction for a mean of 63.1% of genes for ileal CD8 memory T cells and 68.4% for ileal memory B cells, and reached 80 to 90% in the two highest-scoring donors of each pair. For ileal memory B cells, this was higher than scGen (59.6%) and scPRAM (53.9%). For ileal CD8 memory T cells, the three methods were close (63.4% for scGen and 60.5% for scPRAM).

### scXOrGAN translates blood cell expressions into BALF cell profiles of COVID-19 patients

We next tested scXOrGAN on a COVID-19 atlas^16^ with blood and bronchoalveolar lavage fluid (BALF) from the same patients. Eight patients had both samples, two with mild and six with severe disease. We used six patients for training and held out two patients, P-S086 and P-S088, for testing. We translated five cell-type pairs: CD8 T cells, CD4 T cells, B cells and monocytes in blood into macrophages in BALF, and CD8 T cells in blood into epithelial cells in BALF. We chose these pairs from the correlation and cell-count tables for every blood and BALF cell type in each patient (Supplementary Data 2). The same immune cell type was already highly similar between blood and BALF (for example, *r* 0.92 to 0.95 for CD8 T cells), so we used macrophages and epithelial cells, the BALF cell types that differ from blood cells and had enough cells in every patient, as targets.

Figure 3A shows, for each test patient, the average expression of each highly variable gene in real blood cells against that in real BALF cells of the same patient, with one color per cell-type pair. Before translation, the correlation ranged from 0.54 to 0.79. Monocytes in blood were closest to macrophages in BALF (*r* = 0.74 for P-S086 and 0.79 for P-S088). The lowest correlations were for CD8 T cells in blood against macrophages in BALF in P-S086 (*r* = 0.61) and against epithelial cells in BALF in P-S088 (*r* = 0.54). Many genes expressed in BALF cells were nearly silent in the corresponding blood cells, seen as the band of points along the horizontal axis. Figure 3B shows the same comparison for the BALF cell profiles generated by scXOrGAN . After translation, the correlation was 0.92 to 0.98 across all ten pair-patient combinations, and the genes that were silent in blood moved closer to the diagonal. As in the ileum, this mainly reflects recovery of the target cell identity, since giving every cell the average BALF profile of the training patients reaches a similar level (0.94 to 0.97). Figure 3C shows the distribution of per-gene average expression in real and generated BALF cells for three representative combinations (all ten in Supplementary Figure S1). The generated distribution closely resembled the real one in each case.

**Figure 3:**
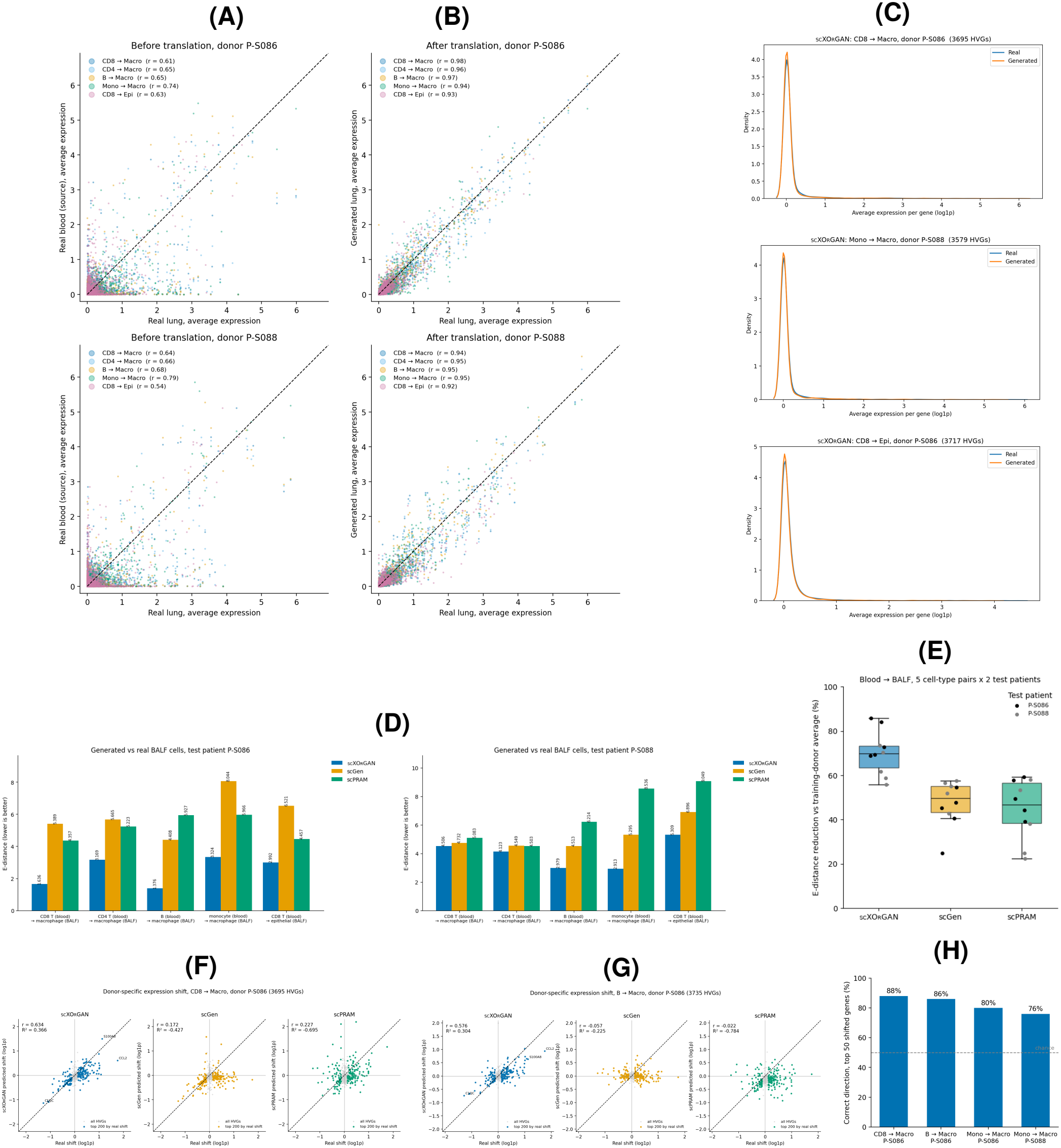
Translation of blood cells into BALF cells by scXOrGAN. (A) Average expression of each highly variable gene in real source cells in blood versus real target cells in BALF, for the two test patients (rows), colored by cell-type pair. (B) The same for scXOrGAN-generated BALF cells versus real BALF cells. *r*, Pearson correlation over the pair’s highly variable genes. (C) Distribution of per-gene average expression in real and generated BALF cells for three representative pair-patient combinations: CD8 T cells in blood to macrophages in BALF (P-S086), monocytes in blood to macrophages in BALF (P-S088), and CD8 T cells in blood to epithelial cells in BALF (P-S086). (D) E-distance between generated and real BALF cells for each cell-type pair and method, for test patients P-S086 (left) and P-S088 (right); lower is better. (E) E-distance reduction relative to the training-donor average (the average training-patient BALF profile assigned to every generated cell), for five pairs and two test patients (*n* = 10 per method). 0% is no better than the training-donor average. Boxes show median and interquartile range; points are individual pair-patient combinations. (F and G) Real versus predicted per-gene shift from the training-patient average for patient P-S086, for CD8 T cells (F) and B cells (G) in blood translated to macrophages in BALF. Colored points are the 200 genes with the largest real shift; labelled genes are the three largest. Dashed line, *y* = *x*. (H) Percentage of the 50 most-shifted genes for which scXOrGAN predicted the correct direction of shift, for four pair-patient combinations; dashed line, chance (50%).

Figure 3D shows the E-distance between generated and real BALF cells for each cell-type pair and method, one panel per test patient. scXOrGAN had the lowest E-distance of the three methods in all ten pair-patient combinations. Figure 3E shows the reduction in E-distance relative to the training-donor average, over all ten combinations. scXOrGAN reduced the E-distance by a median of 69.9% (range 55.9 to 85.9%), compared with 49.8% (24.8 to 57.7%) for scGen and 46.8% (22.3 to 59.4%) for scPRAM.

We also checked whether the generated cell profiles vary from cell to cell as real ones do. For each gene, we compared the variance across cells and the fraction of cells with zero expression between generated and real BALF cells (Supplementary Data 3; one example in Supplementary Figure S2). Across the ten pair-patient combinations, the total per-gene variance of the generated cells, relative to that of real cells, had a median of 0.83 (range 0.64 to 1.25) for SCX-ORGAN, against 0.15 (0.06 to 0.21) for scGen and 0.41 (0.26 to 0.91) for scPRAM, where 1 means the same overall spread. The per-gene fraction of zeros in scXOrGAN -generated cells correlated with that in real cells at *r* = 0.87 to 0.97, higher than for scGen and scPRAM in all ten combinations.

Finally, we explored what is specific to each patient. Figures 3F and 3G plot, for P-S086, the real shift of each gene from the training-patient average against the shift predicted by each method, for CD8 T cells (F) and B cells (G) in blood translated to macrophages in BALF. The shifts predicted by scXOrGAN correlated with the real ones at *r* = 0.63 and 0.58, against *−*0.06 to 0.23 for scGen and scPRAM. Figure 3H shows the percentage of the 50 most-shifted genes for which scXOrGAN predicted the correct direction, for CD8 T cells and B cells to macrophages in P-S086 and for monocytes to macrophages in both patients. In these four combinations, scXOrGAN predicted the correct direction for 76 to 88% of genes, against 50% by chance.

### Generated BALF macrophage profiles represent COVID-19-associated genes

Next, we analyzed whether the generated cell expressions also reproduce the expression of genes linked to COVID-19. We used the genes reported by Liao et al.^24^, who studied macrophages in BALF from patients with moderate or severe COVID-19 and from healthy people. In their data, *FCN1, SPP1* and the inflammatory genes *IL1B, IL6, TNF, CCL2, CCL3, CCL4* and *CCL7* were expressed at high levels in severe disease, and *FABP4* was expressed at high levels in healthy people and in moderate disease. *CXCL9, CXCL10* and *CXCL11* were upregulated in COVID-19 compared with healthy people, and *CXCL16* was expressed at higher levels in moderate than in severe disease. We also added *S100A8, CD14, APOC1* and *MARCO*, which they used as markers of macrophage subsets. This gives 18 genes. A gene is shown only if it is among the highly variable genes of that cell-type pair, so *CXCL16* is missing for CD8 T cells and *CD14* is missing for monocytes.

Figure 4 shows two examples: CD8 T cells in blood translated to macrophages in BALF for patient P-S086 (A), and monocytes in blood translated to macrophages in BALF for patient P-S088 (B). Each plot has three rows: the real blood cells, the generated BALF macrophages and the real BALF macrophages. Dot size shows the share of cells that express the gene, and color shows the mean expression.

**Figure 4:**
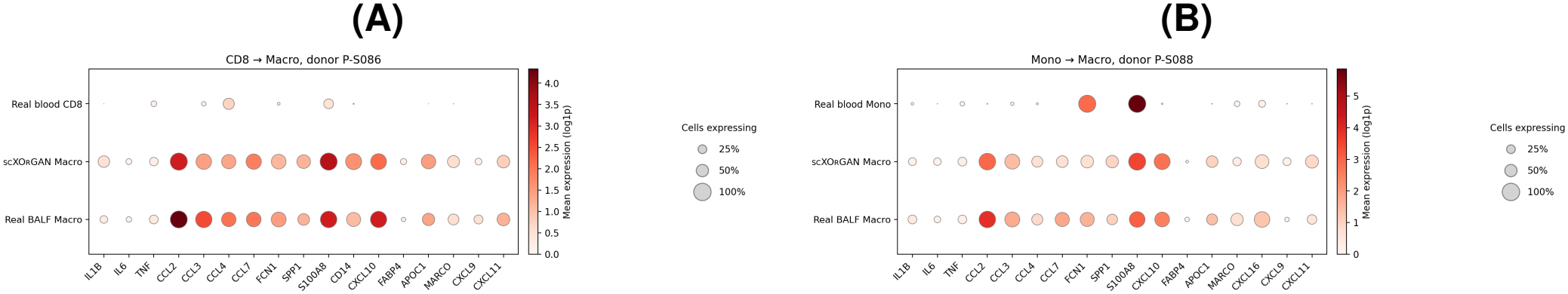
COVID-19-associated genes in generated BALF macrophages. Mean expression (color) and share of cells that express the gene (dot size) for COVID-19 genes from Liao et al. ^24^, in real blood cells (top row), SCX-ORGAN-generated BALF macrophages (middle row) and real BALF macrophages (bottom row). (A) CD8 T cells in blood translated to macrophages in BALF, patient P-S086. (B) Monocytes in blood translated to macrophages in BALF, patient P-S088. Genes that are not among the highly variable genes of the pair are not shown (*CXCL16* in A, *CD14* in B).

Blood CD8 T cells expressed almost none of these genes. The generated BALF macrophage profiles gained expression of most of them. Most of the genes reached levels close to those in real BALF macrophages, while some remained lower than in real BALF macrophages. Blood monocytes were different, because they already had high *FCN1* and *S100A8*. In the generated BALF macrophages, expression of both genes decreased, closer to the real BALF macrophages. So the model changed the source cells instead of copying them. The other six pair-patient combinations are shown in Supplementary Figure S3.

### scXOrGAN translates expressions from blood cells into heart cell profiles of heart failure patients

Finally, we tested scXOrGAN on a heart failure atlas^17^, with scRNA-seq data from blood and heart tissue from the same people. We used data from six individuals: one without heart failure (N-1), two with heart failure caused by a weak and enlarged heart muscle (dilated cardiomyopathy; DCM-2 and DCM-3), and three with heart failure caused by blocked heart arteries (ischemic cardiomyopathy; ICM-1 to ICM-3). We held out DCM-3 and ICM-3 for testing and used the other four people for training. We translated T cells and B cells in blood into fibroblasts and endothelial cells in the heart, which gives four cell-type pairs. The same immune cell type was already highly similar between blood and heart (*r* above 0.9 for B cells and T cells; Supplementary Data 4), so we used non-immune heart cells with enough cells as targets.

Figure 5A shows, for each test patient, the average expression of each gene in real blood cells against that in real heart cells of the same patient, with one color per cell-type pair. Before translation, the correlation was between 0.62 and 0.68 for all four pairs in both patients, and many genes expressed in heart cells were close to zero in blood cells. Figure 5B shows the same comparison for the heart cells generated by scXOrGAN . After translation, most points moved close to the diagonal, and the correlation rose to between 0.90 and 0.97. The highest was for B cells in blood to fibroblasts in the heart in DCM-3 (*r* = 0.97).

**Figure 5:**
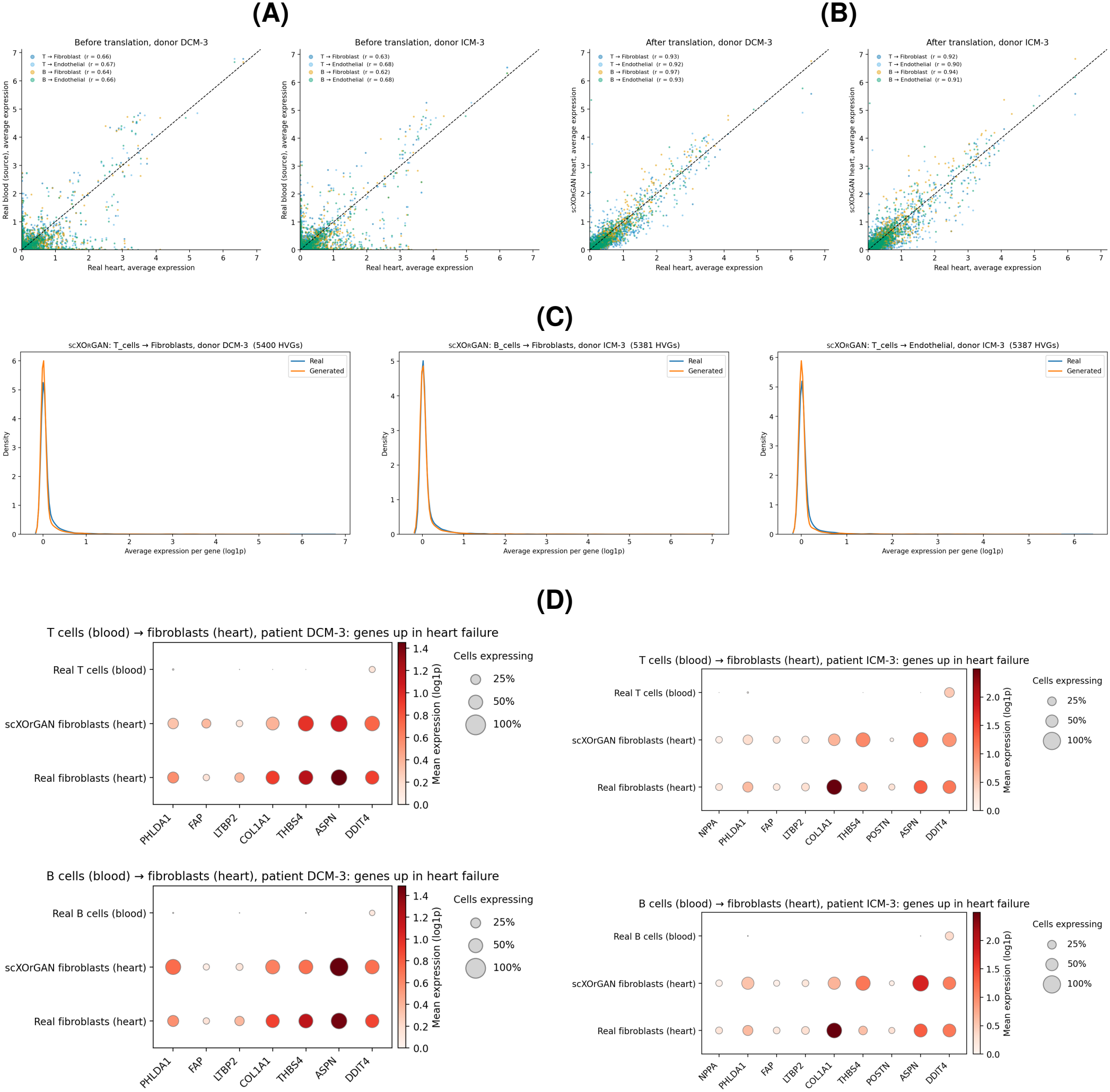
Translation of blood cells into heart cells by scXOrGAN. (A) Average expression of each gene in real source cells in blood versus real target cells in the heart, for test patients DCM-3 (left) and ICM-3 (right), colored by cell-type pair: T cells or B cells in blood to fibroblasts or endothelial cells in the heart. (B) The same for scXOrGAN-generated heart cells versus real heart cells. *r*, Pearson correlation over the pair’s genes; dashed line, *y* = *x*. (C) Distribution of per-gene average expression in real and generated heart cells for three examples: T cells in blood to fibroblasts in the heart (DCM-3, left), B cells in blood to fibroblasts in the heart (ICM-3, middle), and T cells in blood to endothelial cells in the heart (ICM-3, right). (D) Expression of genes upregulated in heart failure in real source cells in blood, scXOrGAN-generated fibroblasts in the heart and real fibroblasts in the heart of the same patient, for T cells (top) and B cells (bottom) in blood, in DCM-3 (left) and ICM-3 (right). Genes are from the 20 most upregulated in heart failure among those significant in bulk RNA sequencing of the left ventricle ^17^, shown if expressed in at least 10% of real fibroblasts of that patient (Methods). Dot size, share of cells expressing the gene; color, mean expression (log1p).

Figure 5C shows the distribution of per-gene average expression in real and generated heart cells for three examples (all eight pair-patient combinations are in Supplementary Figure S4). The generated curves closely matched the real ones. For T cells, the generated cells had slightly more genes close to zero than the real heart cells.

We then asked whether the generated heart cells express genes linked to heart failure. The single-cell data include only one person without heart failure, which is too few to test for differential expression between groups of people. We therefore used the bulk RNA sequencing of the left ventricle from the same study (52 samples with heart failure and 15 without), in which 1,910 genes were differentially expressed after Bonferroni correction. From these, we took the 20 most upregulated genes in heart failure, and for each pair and patient we show those expressed in at least 10% of the real target cells. Figure 5D shows these genes for fibroblasts in the heart. Real T cells and B cells in blood showed little or no expression of them. The fibroblasts generated by scXOrGAN expressed them at levels and in fractions of cells close to those of real fibroblasts in the same patient, including *ASPN, THBS4, LTBP2* and *FAP*. The largest difference was in *COL1A1* in ICM-3, whose expression was lower in the generated fibroblasts than in the real ones.

## CONCLUSION

In this work we asked the question: given the gene expression of single cells from one organ, can we generate the gene expression of single cells from another organ of the same person? We presented scXOrGAN, a model that answers this at single-cell resolution, even though no cell in one organ has a matching cell in the other. scXOrGAN learns from cells that are matched only at the level of the person, using two autoencoders with a shared latent space, cycle consistency, adversarial training and distribution matching.

We tested scXOrGAN on three pairs of organs. In every case, the source cells expressions were far from the target cell ones before translation, and the generated cell profiles were close to the real target profiles after translation. The generated expressions also varied from cell to cell similarly to real cells, instead of collapsing to one average profile. scXOrGAN matched the real cell populations better than two widely used relevant tools, scGen and scPRAM, in almost every test donor of the ileum and in all ten blood-to-BALF combinations. The generated profiles also represented genes linked to diseases. Generated BALF macrophage profiles acquired expression of COVID-19 genes that were almost absent in blood cells, and generated heart fibroblast ones expressed genes that change in patients with heart failures.

Our study nevertheless has several limitations. The BALF and heart datasets have data from only a few patients, so we could test on only two patients each. A large part of the agreement between generated and real cells comes from getting the cell type right, and predicting what is special about each patient remains hard. Larger datasets with more people and paired samples from both organs will be needed to improve this part. Even so, our results show that cross-organ translation at single-cell resolution is possible. With more data, models like scXOrGAN could one day help follow the state of hard-to-reach organs from a simple blood sample.

## METHODS

### Data preprocessing

For each donor we take scRNA-seq data from the cells of one cell type from the source organ and the cells of one cell type from the target organ. The two sides are unpaired at the cell level, so we treat them as two sets of cells that share a donor. Raw counts are normalized to 10,000 counts per cell and then transformed with log(1 + *x*) where *x* is the normalized count.

We then select highly variable genes separately for the source and the target cell type, taking the 2,000 most variable genes on each side using the training donors only, and keep the union of the two lists. For the heart dataset, we also add the 1,910 genes differentially expressed between hearts with and without heart failure (Bonferroni-adjusted *p <* 0.05) in the bulk RNA-seq data of the same study^17^. The same gene list is used for both organs and for every donor of a pair, so no information from the test donors is used to choose genes.

### Model architecture

scXOrGAN consists of two variational autoencoders^19^ that share one latent space, together with three discriminators (Figure 1B). We write *x* for a source-organ expression vector and *y* for a target-organ expression vector. The source autoencoder has encoder *E*_*X*_ and decoder *D*_*X*_, and the target autoencoder has encoder *E*_*Y*_ and decoder *D*_*Y*_ . Each encoder has three fully connected layers of 512, 256 and 128 units, each followed by layer normalization and a ReLU activation, and ends in two linear heads that give the mean and log-variance of a 64-dimensional latent distribution:

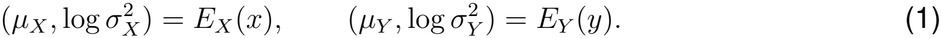

Latent variables are drawn with the reparameterization trick, *z* = *µ* + *σ ⊙ ϵ* with *ϵ ∼ N* (0, *I*). Each decoder mirrors its encoder (128, 256 and 512 units) and ends in a linear layer followed by a ReLU, so that the model cannot produce negative expression values.

Translation is done by cross-wiring the two branches: a cell is encoded by its own encoder and decoded by the other organ’s decoder,

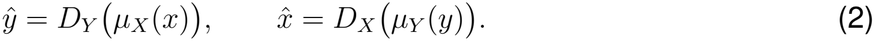

The three discriminators each have one hidden layer of 128 units with a ReLU activation and a single output score. The latent discriminator *D*_*z*_ receives sampled latent vectors and tries to tell source latents from target latents, which pushes the two encoders towards a common latent space. The expression discriminators *D*_*x*_ and *D*_*y*_ receive gene expression vectors and try to tell real cells of their organ from translated ones.

### Loss functions

#### Reconstruction and KL divergence

Each autoencoder rebuilds its own input from a sampled latent. Squared error is summed over genes and averaged over cells, and the latent distributions are pulled towards a standard normal prior *p*(*z*) = *N* (0, *I*):

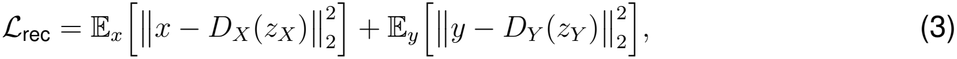

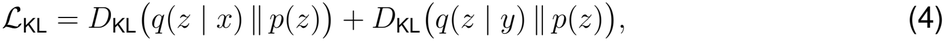

and *ℒ*_VAE_ = *ℒ*_rec_ + *ℒ*_KL_.

#### Cycle consistency

A cell is translated to the other organ and back, and we penalize the deviation from the original position.^9^:

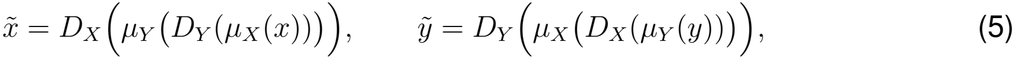

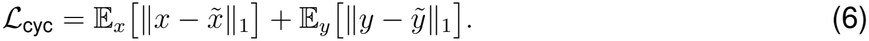

Because the two organs are unpaired at the cell level, this is the term that ties a translated cell to the cell it came from.

#### Distribution matching

The maximum mean discrepancy (MMD)^21^ compares the *m* generated target cells *ŷ* with the *n* real target cells *y* of the same donor in the current step, in gene expression space:

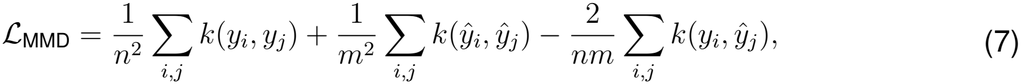

with a multi-scale Gaussian kernel

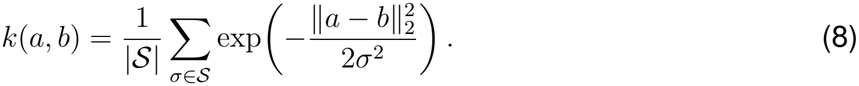

The bandwidths are *S* = *{*0.5, 1, 2, 4*} × δ*, where *δ* is the median distance between up to 1,000 real target cells of the training donors. They are set once before training and kept fixed. Reconstruction and cycle consistency act on single cells; MMD is the only term that compares whole groups of cells.

#### Adversarial terms

Each discriminator *D* is trained with binary cross-entropy on its scores, and the encoders and decoders are trained against it with the generator loss,

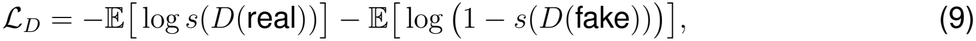

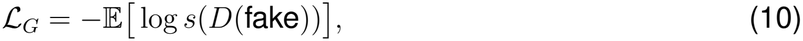

where *s*(*·*) is the logistic function^20^. The adversarial term *ℒ*_adv_ is the sum of the generator losses for *D*_*z*_, *D*_*x*_ and *D*_*y*_.

#### Total objective

The model is trained to minimize

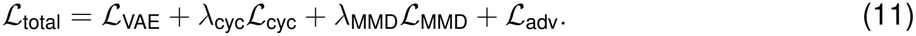

Before training, we measure each term on one batch and set *λ*_cyc_ and *λ*_MMD_ so that the cycle and MMD terms each start at 10% of the reconstruction term. The terms differ by orders of magnitude in their natural scale, so this keeps every loss on a similar scale and stops any one term from dominating.

### Training and inference

Source and target cells are unpaired at the cell level but matched at the donor level. We use this with donor-wise batching: every training step takes 32 source cells and 32 target cells from a single training donor *P*_*i*_,

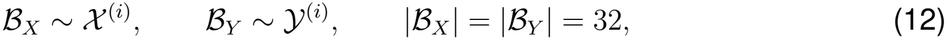

where *X* ^(*i*)^ and *Y*^(*i*)^ are the source and target cells of *P*_*i*_. This keeps the donor fixed within a step, so the MMD term and the expression discriminators always compare generated cells with real cells of the same person.

One epoch is a full pass over all cells of both organs for every training donor. In each step, the three discriminators are updated first, each on its own loss, and then the encoders and decoders are updated once on the total objective. All parameters are optimized with Adam at a learning rate of 10^*−*4^ for 1,000 epochs, with random seed 0, and the model after the last epoch is used. At test time, every source-organ cell of a test donor is encoded with *E*_*X*_ and its posterior mean is decoded with *D*_*Y*_, giving one generated target-organ cell per input cell. No target-organ data from a test donor are used at any stage.

### Evaluation

We compare generated and real target cell profiles of each test donor as groups, since the two organs have no matching cells. For average expression, we take the mean of each gene over the generated and over the real cells, and compute the Pearson correlation of these two profiles over the pair’s genes. For distributions, we use the E-distance^22,23^, 2 E∥*a − b*∥ *−* E∥*a − a*^*′*^∥ *−* E∥*b − b*^*′*^∥, with plain Euclidean distances. It is computed in a 50-dimensional principal component space fit on real target cells of all donors (at most 2,000 cells per donor), on equal numbers of real and generated cells (at most 1,500), and averaged over three random draws. As a reference, we used the training-donor average, in which the mean expression profile of the target cell type in the training donors, with each donor weighted equally, is assigned to every cell. We report each method’s reduction in E-distance relative to this reference, 100 (1 *− E*_method_*/E*_average_).

To test what is specific to a donor, we compute the shift of each gene as the donor’s mean expression minus the training-donor average, for both real and generated cell profiles. We report the Pearson correlation between real and predicted shifts, and the percentage of the 50 genes with the largest real shift for which the predicted shift has the correct sign. In the dot plots, dot size is the share of cells with expression above zero and color is the mean log(1 + *x*) expression.

### Heart failure differentially expressed gene selection

The single-cell heart data include only one person without heart failure (N-1), so differential expression cannot be tested between groups of people at the single-cell level. We therefore used the bulk RNA sequencing of the left ventricle from the same study^17^: 52 samples from patients with heart failure and 15 samples without heart failure. The bulk samples carry no label for the cause of heart failure, so the comparison is heart failure against no heart failure.

We tested differential expression with DESeq2 (version 1.52.0)^25^ in R (version 4.6.1), with heart failure status as the only factor. We applied Bonferroni correction over all 42,718 genes with a p-value, and 1,910 genes were significant (adjusted *p <* 0.05).

From these 1,910 genes, the 20 most upregulated in heart failure were selected with the Biomni AI agent^26^. Genes were ranked by fold change within the significant set. Long non-coding RNAs, pseudogenes, unnamed loci and genes that mark tissues other than the heart (for example, salivary, pancreatic or neuronal genes) were removed, and the remaining genes were checked against known heart failure biology.

Because bulk left ventricle is mostly cardiomyocytes, some of these genes are not expressed in fibroblasts or endothelial cells. For each cell-type pair and test patient, we therefore show a gene only if it is in the pair’s gene set, since scXOrGAN generates only those genes, and if it is expressed (count above zero) in at least 10% of the real target cells of that patient.

## RESOURCE AVAILABILITY

### Lead contact

Requests for further information and resources should be directed to and will be fulfilled by the lead contacts, MAHS and AR.

### Data availability

All data are public and no new data were generated. IBDverse is available from CELLxGENE at https://cellxgene.cziscience.com/collections/7f7fdf50-aa0e-4a6f-979c-53655cac9d5d (dataset 0e5c75da-7bc9-4a2c-83fc-ef1e378bd5f9). GSE158055 and GSE145154 are available from the NCBI Gene Expression Omnibus at https://www.ncbi.nlm.nih.gov/geo/query/acc.cgi?acc=GSE158055 and https://www.ncbi.nlm.nih.gov/geo/query/acc.cgi?acc=GSE145154. The method is not specific to any of these datasets. It needs only two sets of cells from two organs of the same donors, with cell type labels.

