## Supplementary Information for "scXOrGAN: A Generative Framework for Cross-Organ Single-Cell Transcriptomic Translation"

### Supplementary Data

**Supplementary Data 1** (`Supplementary_Data_1.xlsx`). Pearson correlation between the average expression of each cell type in the ileum and each cell type in the rectum, for the healthy donors of IBDverse with at least 200 cells in both tissues (209 donors), computed on all 31,196 genes detected in at least one cell (counts per 10,000, log1p). A pair is scored in a donor only if the donor has at least 100 source cells in the ileum and 100 target cells in the rectum. Sheets give the median correlation across donors for every pair, the number of qualifying donors for every pair, a list of all pairs with their number of donors and the median, mean, minimum and maximum correlation, and the correlation in each donor. These tables were used to choose the cell-type pairs for translation.

**Supplementary Data 2** (`Supplementary_Data_2.xlsx`). Pearson correlation between the average expression of each cell type in the blood and each cell type in the BALF, for each of the eight patients in the COVID-19 atlas with both blood and BALF samples, computed on all genes before any training. Below each correlation table is the number of cells in the smaller of the two groups for each pair. One sheet per patient. Blank cells mark pairs with fewer than 20 cells in either group. These tables were used to choose the cell-type pairs for translation.

**Supplementary Data 3** (`Supplementary_Data_3.xlsx`). Cell-to-cell variation of real and generated cells in the bronchoalveolar lavage fluid (BALF), for the ten test donor and cell-type pairs and for scXOrGAN, scGen and scPRAM. For each gene, the variance across cells and the fraction of cells with zero expression were computed in real and in generated cells (log-normalized expression, highly variable genes). Columns give the Pearson correlation between real and generated per-gene variance, the ratio of total generated to total real variance (1 means the same spread), the Pearson correlation between real and generated per-gene zero fraction, and the average zero fraction of real and of generated cells.

**Supplementary Data 4** (`Supplementary_Data_4.xlsx`). Pearson correlation between the average expression of each cell type in the blood and each cell type in the heart, for every patient in the heart failure atlas, computed on all genes detected in at least one cell. Below each correlation table is the number of cells in the smaller of the two groups for each pair. One sheet per patient. These tables were used to choose the cell-type pairs for translation.

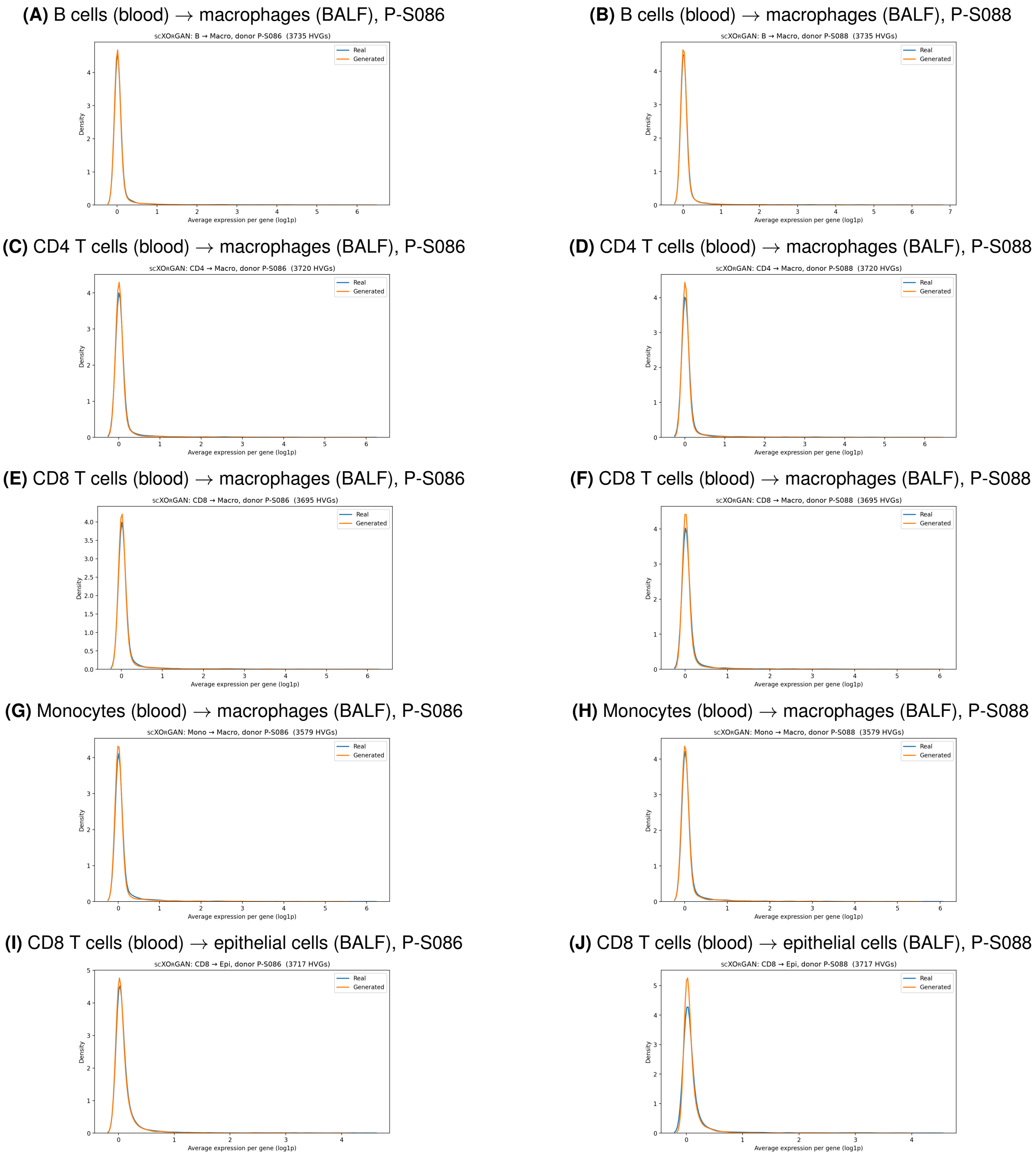


**Figure S1: Distribution of per-gene average expression in real and scXOrGAN-generated cells in the BALF, for all ten test donor and cell-type pairs.** Each panel shows one translation from the blood to the BALF for one test donor (P-S086 or P-S088): B cells (A and B), CD4 T cells (C and D), CD8 T cells (E and F) and monocytes (G and H) in the blood to macrophages in the BALF, and CD8 T cells in the blood to epithelial cells in the BALF (I and J). Related to Figure 3.

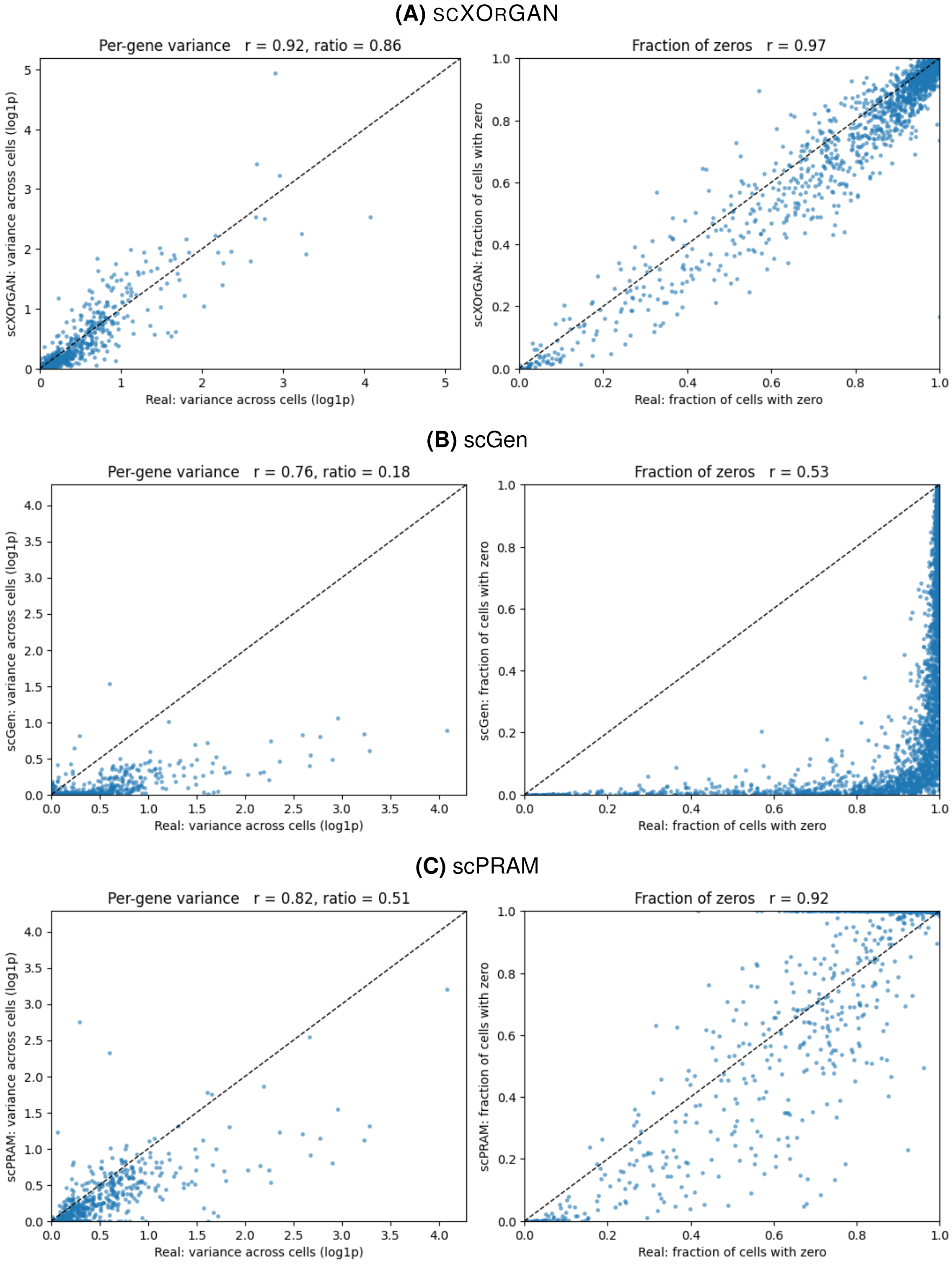


**Figure S2: Cell-to-cell variation of generated macrophages in the BALF, translated from CD8 T cells in the blood, for test donor P-S086.** Generated cells from scXOrGAN (A), scGen (B) and scPRAM (C) compared with real macrophages in the BALF, over 3,695 highly variable genes. Left, variance of each gene across cells (log1p); right, fraction of cells with zero expression for each gene. Each point is a gene; dashed line, $y = x$. $r$, Pearson correlation; ratio, generated to real variance. Values for all ten test donor and cell-type pairs are in Supplementary Data 3.

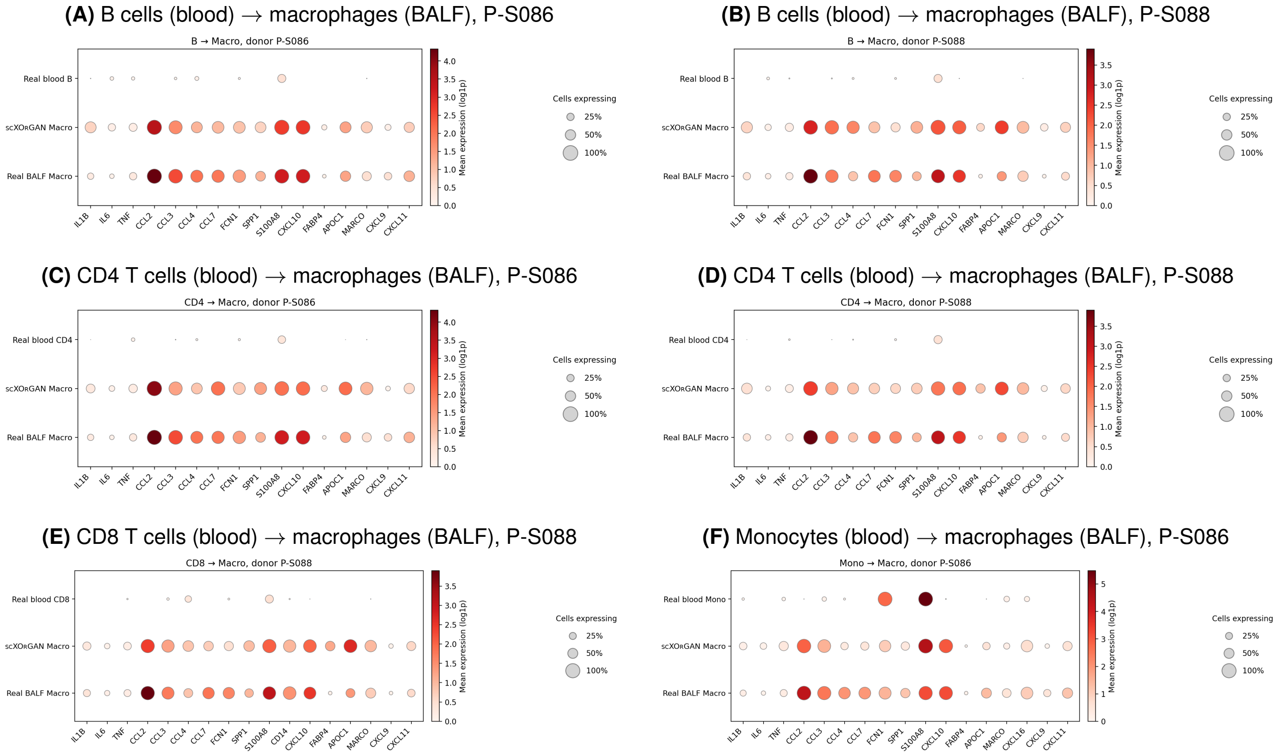


**Figure S3: Expression of COVID-19-associated genes in real and scXOrGAN-generated macrophages in the BALF, for the six donor and cell-type pairs not shown in Figure 4.** Translations from B cells (A and B) and CD4 T cells (C and D) in the blood for test donors P-S086 and P-S088, from CD8 T cells in the blood for P-S088 (E), and from monocytes in the blood for P-S086 (F), all to macrophages in the BALF. Dot size shows the fraction of cells expressing the gene; color shows average expression. Genes are from Liao et al. (2020). Related to Figure 4.

**(A)** B cells (blood) → endothelial cells (heart), DCM-3

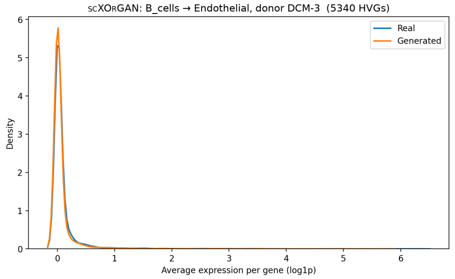


**(B)** B cells (blood) → endothelial cells (heart), ICM-3

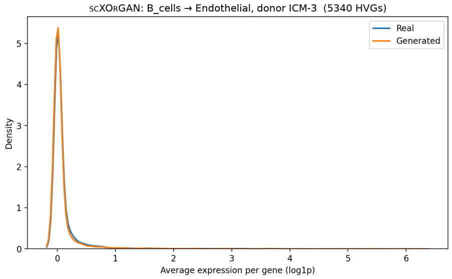


**(C)** B cells (blood) → fibroblasts (heart), DCM-3

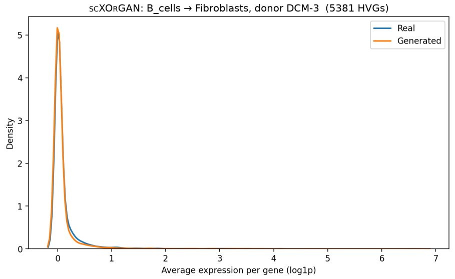


**(D)** B cells (blood) → fibroblasts (heart), ICM-3

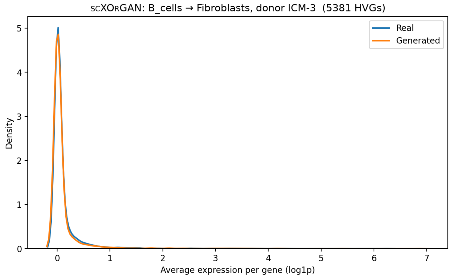


**(E)** T cells (blood) → endothelial cells (heart), DCM-3

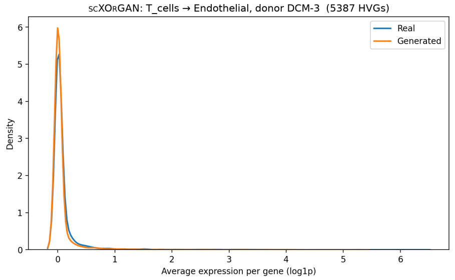


**(F)** T cells (blood) → endothelial cells (heart), ICM-3

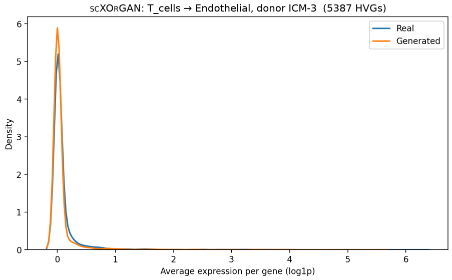


**(G)** T cells (blood) → fibroblasts (heart), DCM-3

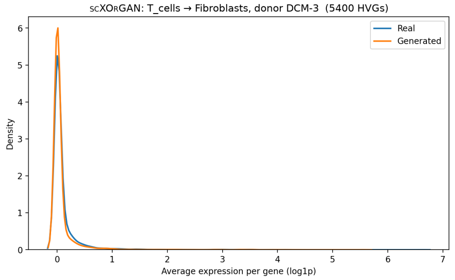


**(H)** T cells (blood) → fibroblasts (heart), ICM-3

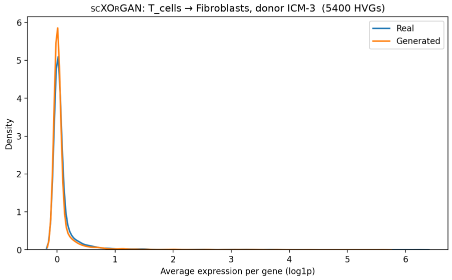


**Figure S4: Distribution of per-gene average expression in real and scXOrGAN-generated cells in the heart.** Each panel shows one translation from the blood to the heart for one test donor (DCM-3 or ICM-3): B cells (A–D) and T cells (E–H) in the blood to endothelial cells and fibroblasts in the heart. Related to Figure 5.
